# Overcoming the speed limit of four‐way DNA branch migration with bulges in toeholds

**DOI:** 10.1101/2023.05.15.540824

**Authors:** Francesca Smith, Aditya Sengar, Guy‐Bart V. Stan, Thomas E. Ouldridge, Molly Stevens, John Goertz, Wooli Bae

**Affiliations:** Department of Physics, Faculty of Engineering and Physical Sciences, University of Surrey, GuildfordGU2 7XH, U.K.; Imperial College Centre of Excellence in Synthetic Biology and Department of Bioengineering, Imperial College London, South Kensington Campus, London SW7 2AZ, U.K.; Department of Materials, Department of Bioengineering and Institute of Biomedical Engineering, Imperial College London, London, SW7 2AZ, U.K.

## Abstract

Dynamic DNA nanotechnology involves the use of DNA strands to create programmable reaction networks and nanodevices. The key reaction in dynamic DNA nanotechnology is the exchange of DNA strands between different molecular species, which is achieved through three-way and four-way DNA exchange reactions. While both of these reactions have been widely used to build reaction circuits, the four-way exchange reaction has traditionally been slower and less efficient than the three-way reaction. In this paper, we describe a new mechanism to optimise the kinetics of the four-way DNA exchange reaction by adding bulges to the toeholds of the four-way DNA complexes involved in the reaction. These bulges facilitate an alternative branch migration mechanism and destabilise the four-way DNA junction, increasing the branch migration rate and unbinding rate of the four-way exchange reaction, bringing it closer to the kinetics of the three-way reaction. This new mechanism has the potential to expand the field of dynamic DNA nanotechnology by enabling efficient four-way DNA exchange reactions for in vivo applications.

## Introduction

The utility of DNA as a building material has progressed significantly within the field of DNA nanotechnology^1^. Meanwhile, The falling cost of nucleic acid synthesis in combination with an improved understanding of nucleic acid assembly and programmability has unlocked our ability to engineer increasingly complex, composable nucleic acid reaction networks in the field of dynamic DNA nanotechnology^2-7^. Applications in the field of dynamic DNA nanotechnology include dynamic nanomachines^8-10^, sensing devices^11-12^ and molecular computing reaction networks^13-17^.

Toehold-mediated strand displacement (TMSD), in which DNA and RNA strands are exchanged in a highly specific manner with controllable kinetics^18-19^, is the fundamental reaction in this field. A typical TMSD reaction involves the exchange of one strand for another within a complex via three-way DNA branch migration(Figure 1a). In this reaction, a single-stranded invader hybridises to a substrate-incumbent complex via a short single-stranded overhang, known as the toehold. Following toehold binding, a three-way junction forms between the invader, substrate and incumbent strands which can migrate back and forth in a random walk fashion until the invader either completely detaches or displaces the incumbent strand.

**Figure 1.**
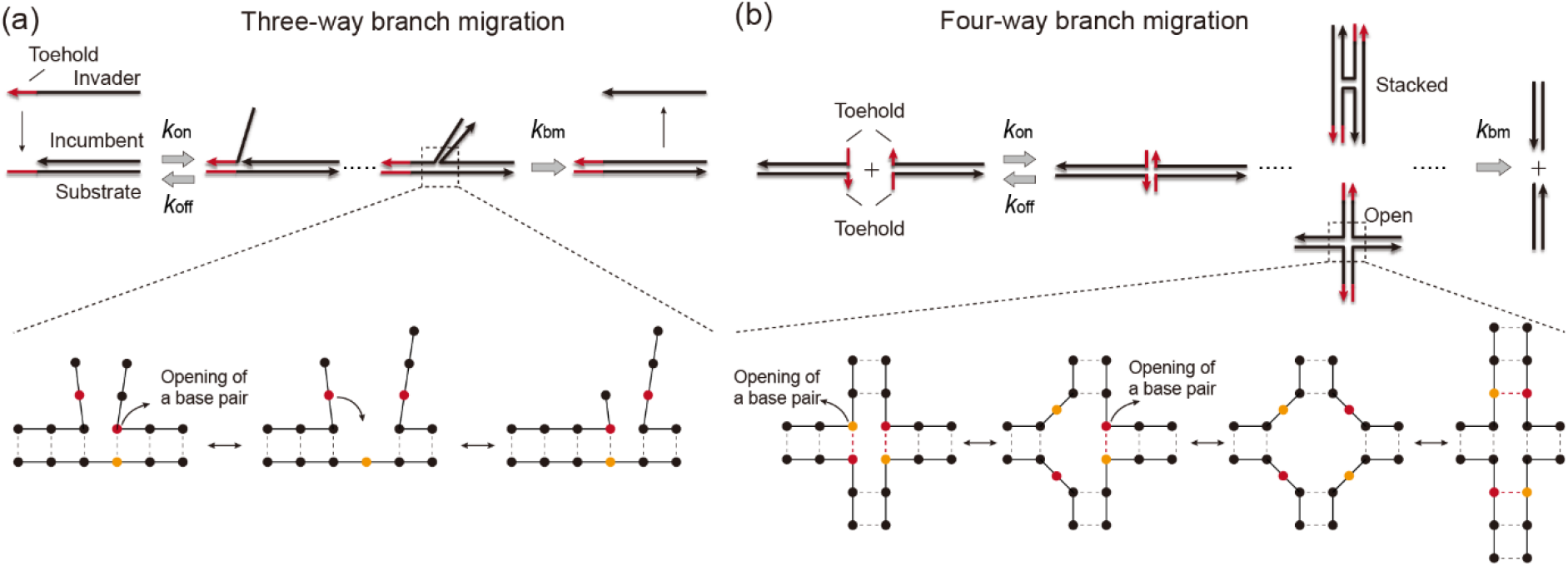
Comparison of three-way and four-way branch migration mechanisms. a) In three-way branch migration, a single-stranded invader hybridises to an incumbent-substrate complex via the toehold before progressively displacing the incumbent strand (upper scheme). During the displacement, branch migration is thought to occur^19, 23^ by stochastic opening of an incumbent-substrate base-pair and its replacement by an invader-substrate base-pair (lower scheme). b) In four-way branch migration, two duplexes hybridise via two toehold sequences, forming a Holliday junction (upper scheme). Here, branch migration involves the exchange of two base-pairs (lower scheme).

Notably, while three-way branch migrations are more frequently employed in dynamic nucleic acid reaction circuits, one can also use a four-way branch migration mechanism in dynamic molecular reactions^20-22^. Four-way branch migration requires the formation of a Holliday junction between two DNA duplexes via hybridisation of two toehold sequences (Figure 1b). The Holliday junction is an important structure that appears during *in vivo* homologous recombination^23^, which is foundational to DNA double-strand break repair^24^ and DNA replication^25^. The crystal structure of the Holliday junction revealed that the four-arms of the Holliday junctions form a stable stacked monomer in the presence of high salt concentration (Figure 1b, stacked form) which is interchangeable with its open form^26-27^. Branch migration can proceed by the simultaneous exchange of two basepairs at the junction, until either complete dissociation into the original duplexes or exchange of strands is complete^23^. The four-way branch migration scheme holds a number of advantages over three-way branch while keeping the ability to design complex reaction circuits. The requirement for hybridisation of two toehold sequences generally increases the specificity and the number of orthogonal species in strands interaction. Also, four-way branch migration eliminates the need for single-stranded reaction species and so minimises the potential issues associated with secondary self-complementarity or undesired cross-talk enabling less noisy designs especially in living systems due to higher stability^28-29^.

One of the main reasons for the success of three-way branch migration is its reaction kinetics. A strand displacement reaction is often modelled as a two-step process represented by three reaction rates *k*_on_, *k*_off_ and *k*_bm_, which are the rate constants of bimolecular binding through toehold, unimolecular unbinding and branch migration, respectively (Figure 1a and b)^19^. In a three-way branch migration reaction, *k*_bm_, which sets the speed limit of the reaction is typically high^19^. Also, *k*_off_ is high for incorrect toeholds and low for matching toeholds. This makes a three-way branch migration reaction efficient with fast kinetics for correct toeholds and slow kinetics for incorrect toeholds.

However, this is not the case for current implementations of four-way branch migration. The *k*_bm_ of four-way branch migration is two to three orders of magnitude lower that that of three-way branch migration^23^. This difference is due to differences in their molecular mechanisms of the branch migration. Three-way branch migration requires the spontaneous opening of a single base-pair followed by subsequent formation of a new base-pair to achieve a single branch-migration step (Figure 1a). Contrarily, four-way branch migration would rely on the simultaneous opening of two base-pairs followed by subsequent formation of two new base-pairs to achieve a complete branch migration step at the junction (Figure 1b). This effect further enhanced by additional stabilisation of stacked forms of a Holliday junction at high salt (Figure 1b)^27, 30^.

In this work, we introduce a new mechanism for four-way branch migration that increases *k*_bm_ as well as *k*_off_, making the four-way branch migration more efficient and comparable to the three-way branch migration. We use oxDNA simulation^31^ to explain aspects of the system performance. Such a design framework opens up the possibility of constructing rapid, controllable reaction circuits using four-way branch migration in the field of dynamic nucleic acid nanotechnology.

## Results

### Introduction of a single bulge

We hypothesised that the presence of a bulge (unpaired base) at the Holliday junction should be sufficient to trigger an improved four-way branch migration mechanism, with increased reaction kinetics (Figure 2a). A bulge at the junction can minimise the thermodynamic penalty associated with each branch migration step, as the opening of a single base-pair is sufficient for branch migration initiation. Also, the bulge should destabilise the stacked form of the Holliday junction to further facilitate branch migration. To help understanding the mechanism, we visualise the progression of a four-way branch migration in the presence of a bulge in Figure 2a. In state 1, a bulge forms adjacent to the toeholds between arm 1 and arm 3 of the DNA junction, as a result of toehold hybridisation. Thermal fluctuations allow a base-pair in arm 4 to open (state 2). The revealed base *n’* can then bind to the free nucleotide at the bulge (*n*), forming a new base-pair. Overall, this process causes arm 4 to be reduced in length by one base-pair, while arm 1 is extended correspondingly. We refer to this step from state 1 to state 3 as a half-migration step as only two of the arms have been altered. A second half-migration step led by the opening of a base pair in arm 3 (state 4) completes a single branch migration step. At the end of this step (state 5), the junction and the bulge have migrated by one step and the bulge returns to its original position relative to the four-way junction(between the arm 1 and the arm 3).

**Figure 2.**
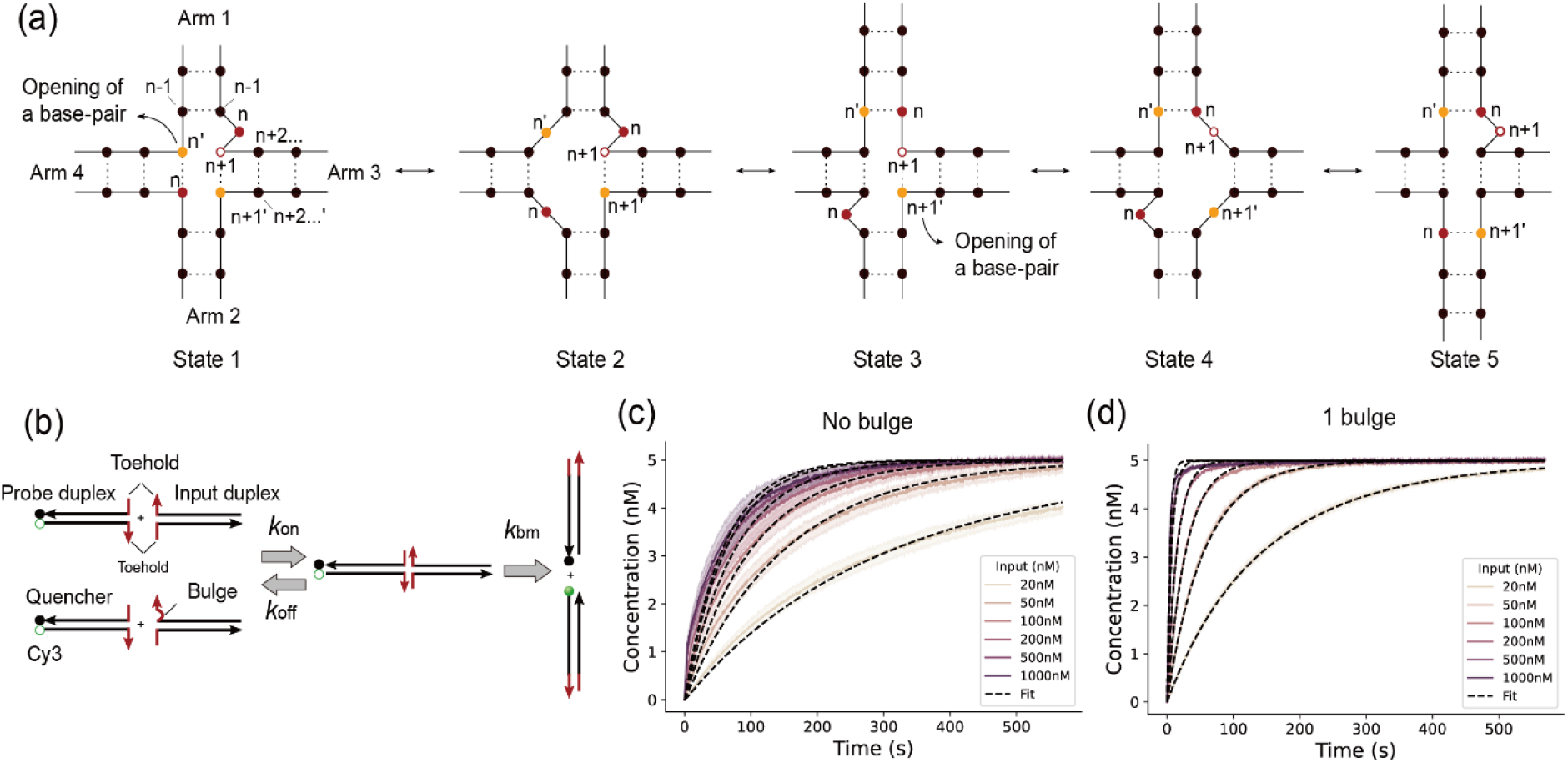
Four-way strand exchange with a single bulge. a) Proposed molecular states during four-way strand exchange with a single bulge. b) The experimental design to compare the four-way branch migration reaction kinetics in the presence and absence of a single bulge. Complete strand displacement recovers a fluorescent product that acts as a reporter of reaction kinetics. Rate constants for our 3-parameter model of four-way branch migration (*k*_on_, *k*_off_ and *k*_bm_) are also indicated. c, d) Normalised fluorescent traces obtained by combining 5 nM of probe duplex and 20 nM – 1000 nM of input duplex in the absence (c) or presence of a single bulge (d). Shaded regions represent standard deviation calculated from three independent experiments. Black, dashed lines represent fitted curves using the three reaction parameters in (b) with only a single set of parameters used for the six different concentrations.

Notably, this reaction mechanism enforces a constraint on the sequence of the branch migration domain as base *n* must be complementary to base *n+1’*, as is evident from state 5. This constraint applies throughout the entire displacement domain. Consequently, the branch migration domain is limited to poly-A or poly-T sequences, as poly-G sequences cannot be synthesised.

In order to assess the effect of introducing a bulge on the kinetics of a four-way branch migration, we employed a simple design involving two DNA duplexes (Figure 2b, Supplementary table 1). Each strand in the first duplex, known as the input duplex, has a 5 nt toehold that is complementary to one toehold on the second duplex, known as the probe duplex. The probe duplex has a fluorophore (Cy3) and a quencher. Completion of strand exchange therefore results in the separation of the quencher and fluorophore and formation of a fluorescent product, which acts as a readout that can be used to estimate the kinetics of this four-way branch migration process. In addition, we designed a blocker strand complementary to the two input strands (with and without bulge) to limit undesired three-way branch migration reactions coming from any single-stranded species that could be present in the solution (See *Materials and methods* for details).

When we mixed different concentrations of input duplex ranging from 20 nM to 1000 nM, with 5 nM of probe duplex, the input duplex with a bulge exhibited much faster strand exchange (Figure 3a and b), particularly at higher concentrations of the input strand. Notably, as the concentration of input strand is increased, input duplexes with a bulge finished strand exchange within seconds while those without a bulge reached a speed limit at hundreds of seconds, regardless of the concentration of input.

**Figure 3.**
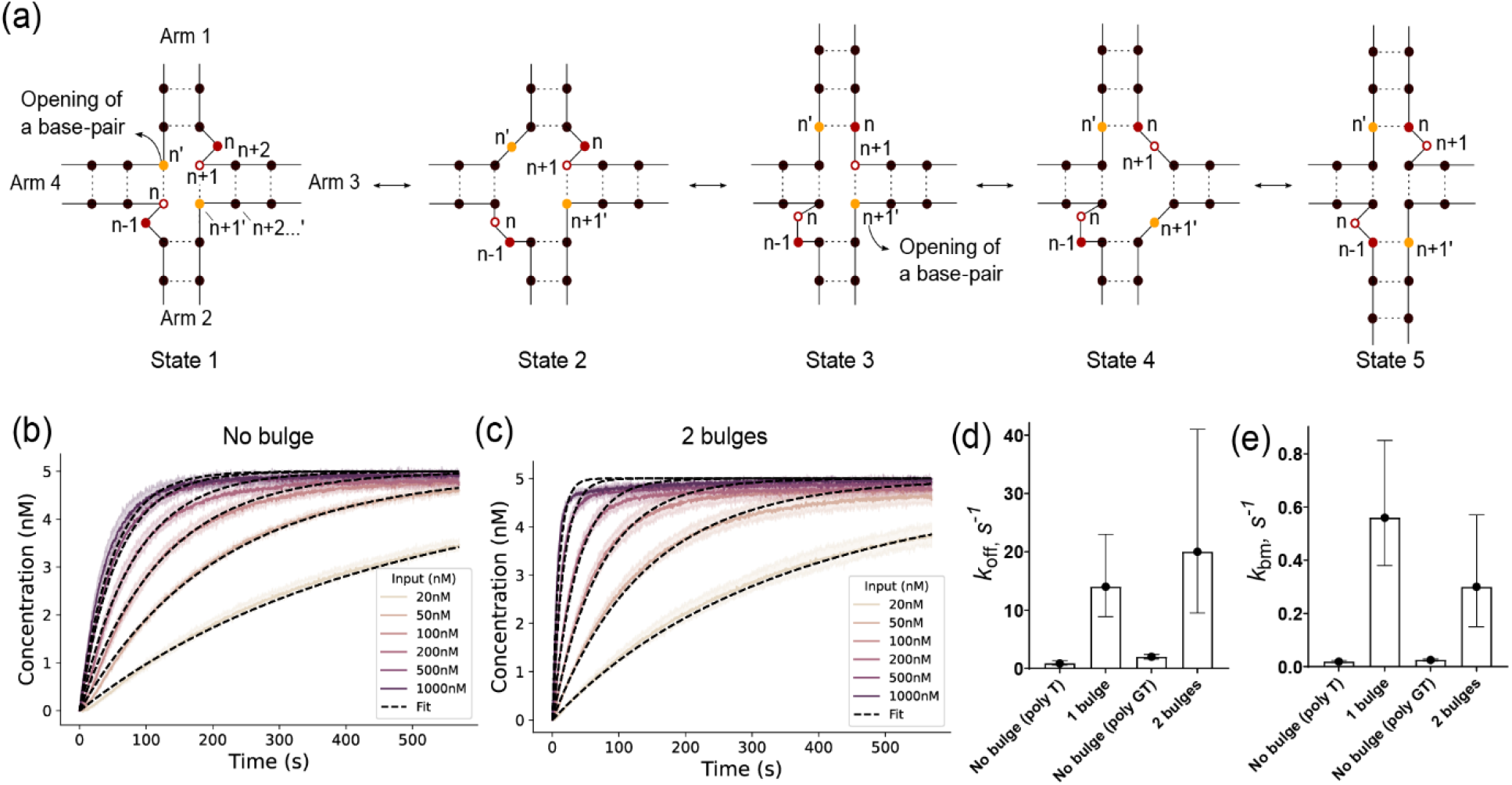
Effect of two bulges on four-way branch migration kinetics. a) Proposed molecular states during four-way strand exchange with a single bulge. In state 1, the two toeholds hybridise to form a bulge between arm 1 and arm 3, and a second bulge between arm 2 and arm 4 at the four-way junction. In state 2, a base-pair opens in arm 4, revealing a base complementary to the bulge nucleotide between arm 1 and arm 3. The binding of the bulge nucleotide to the base n’ forms a new base-pair and generation of a 2 nt bulge between arm 2 and arm 4 (state 3). In state 4, a base-pair opens in arm 2, allowing formation of a new base-pair and formation of two bulges their original position in the junction (state 5). If a base-pair opens in arm 2 initially, this process will occur in the reverse orientation at the junction. b, c) Normalised fluorescent traces from combining 5nM of probe duplex and 20 nM – 1000 nM of input duplex in the absence (b) or presence (c) of two bulges. Shaded regions represent standard deviation calculated from three independent experiments. Black, dashed lines represent fits to the experimental data. d, e) *k*_off_ values (d) and *k*_bm_ values (e) from the fitting (See the Methods for details).

### Introduction of two bulges

We next tested the effect of adding a second bulge at the Holliday junction to see if our mechanism can be generalised and if the speed limit can be further increased by further destabilising the junction. The extension would also relieve the sequence constraint on the branch migration domain to some degree. As in the single bulge case, we hypothesise that the mechanism of four-way branch migration with two bulges involves 5 states (Figure 3a). In state 1, the hybridisation of the toeholds results in formation of a bulge (at base *n*) between arm 1 and arm 3 and a second bulge (at base *n-1*) between arms 2 and 4. Notably, opening of a single base-pair in either arm 2 or arm 3 should be able to initiate a half-migration step. Assuming a base-pair opens in arm 4 (state 2), a new base-pair can form in arm 1 and a 2 nt bulge forms between arm 2 and arm 4 (state 3). We refer to this step as a half-migration step. In state 4 a base-pair opens in arm 3, facilitating formation of a new base-pair with one nucleotide in the 2 nt bulge (state 5). One complete strand displacement step regenerates two bulges at the original positions relative to the four-way junction. Notably, the presence of a second bulge reduces the sequence constraints on the displacement domain as now *n-1* must only be complementary to *n+1’* as is evident in state 5, throughout the displacement domain. As such, in these experiments the displacement domains are composed of poly-GT/poly-CA sequences.

Comparing the kinetics of input duplexes with and without two bulges, we observed once again that the presence of two bulges increased the speed limit of the strand exchange reactions (Figure 3b, c). The reaction in the absence of bulges (Figure 3b) had similar speed to the previous reaction in the absence of bulges for single bulge experiments (Figure 2c). In the presence of two bulges, the strand exchange reactions finished within seconds at high input DNA concentration although they were somewhatt slower than the equivalent reactionsin the presence of a single bulge (Figure 3c).

### Fitting of strand exchange reactions with a 3-parameter model

To further quantify the kinetics of our four-way strand displacement systems, we considered a 3-parameter model to estimate the reaction kinetics. We define *k*_on_, *k*_off_ and *k*_bm_ as the rate constants of bimolecular binding, unimolecular unbinding and branch migration, respectively (Figure 2b). Note that in doing so, we approximate the branch migration process as a single step; this level of detail proved sufficient to describe systems like ours ^19^. As the difference in concentrations of input complex should not affect the reaction rates, we fit all curves in the same panel (Figure 2c, d and Figure 3b, c) with the same set of parameters. However, attempting to fit all three rate constants did not provide reliable rate estimates, since not all parameters were reliably individually constrained by the data (Supplementary note 1). As such, we tried to reduce the parameter space by fixing one of the parameters. We found that fitting *k*_off_ and *k*_bm_ while fixing *k*_on_ to the previously reported maximum rate of hybridization ^32^ gave reasonable values and good fits to all six kinetic curves in each graph (dashed lines in Figure 2c, 2d and 3c, 3d). We confirmed the suitability of our assumption that *k*_on_ = 10^7^ *M*^−1^ *s*^−1^ by systematically altering the fixed *k*_on_ value and evaluating its effect on the *k*_bm_ estimate. Across all experiments, we estimated a similar branch migration rate constant (within 2.5%) for a range of fixed *k*_on_ values from *k*_on_ = 10^6^ *M*^−1^ *s*^−1^ to *k*_on_ = 10^8^ *M*^−1^ *s*^−1^ (Figure S1). For single bulge experiments, *k*_off_ was estimated at 1.4 × 10^1^ *s*^−1^, 95% confidence interval (*CI*) [8.9, 23.0] and 8.6 × 10^−1^ *s*^−1^, 95% *CI* [0.55, 1.3] in the presence and absence of a bulge, respectively (Figure 3d). The mean *k*_bm_ estimate was 5.6 × 10^−1^ *s*^−1^, 95% *CI* [0.38, 0.85] and 1.9 × 10^−2^ *s*^−1^, 95% *CI* [0.015, 0.024] in the presence and absence of a bulge, respectively (Figure 3e). Thus, we concluded that the presence of a bulge accelerated the branch migration process by a factor of approximately 30 times, and increased unbinding of the toeholds holding together the four-way DNA junction by approximately a factor of 20.

We estimated the values of *k*_off_ and *k*_bm_ for two bulges experiments as well. The global fitting protocol provided a mean *k*_off_ estimate of 2.0 × 10^1^ *s*^−1^, 95% *CI* [9.5, 41.0] and 2.0 *s*^−1^, 95% *CI* [1.6, 2.4] in the presence and absence of the two bulges, respectively (Figure 3e). We estimated mean *k*_bm_ values of 3.0 × 10^−1^ *s*^−1^, 95% *CI* [0.15, 0.57] and 2.5 × 10^−2^ *s*^−1^, 95% *CI* [0.021, 0.029] in the presence and absence of the two bulges, respectively (Figure 3f). Similar to the single bulge system, the presence of two bulges destabilised the four-way DNA junction while significantly increasing the branch migration rate.

For both the single and double bulge case, the increase in *k*_bm_ and increase in *k*_off_ have largely compensatory effects when the concentration of probe is low, and binding events are rare. Probes with bulges can perform branch migration faster, but spend less time bound and therefore the probability of successful displacement during a single probe binding event is similar to probes without bulges. Since both bind at a similar rate, the overall reaction rate is similar, as can be seen in Figure 2 and Figure 3. When the probe concentration is high, however, binding events are rapid and so the actual amount of time spent in the four-stranded complex becomes significant in determining reaction kinetics, favouring the designs with bulges in which the resolution of the Holliday junction is much faster. At the highest concentrations, the process becomes effectively first order with a rate entirely determined by *k*_bm_. The systems without bulges clearly show this effect in Figures 2c and 3b; the increasing probe concentration has an ever decreasing return in terms of the reaction rate, tending towards a limiting curve with a relatively low rate set by *k*_bm_. By contrast, the systems with bulges in Figures 2d and 3c exhibit much higher limiting reaction rates. We also note that longer toeholds should have similar effects to high concentrations by increasing the dwell time of the boud states.

### oxDNA simulation

Although the introduction of bulges successfully increased the speed limit of strand exchange reaction and destabilised the Holliday junction, the increase of *k*_bm_ in the presence bulges was not as big as we expected from a picture in which the bulges both reduce the number of base pairs that must be disrupted at anyone time and destabilize the relatively immobile, stacked conformation.

This unexpected behaviour was particularly notable for the double bulge system, which had an estimated *k*_bm_ that was lower than for the single bulge system. We therefore performed oxDNA simulations to further understand the molecular behaviour of our system (Figure 4a)^31, 33^. The simulations were setup and run for comparable experimental conditions: identical temperature, salt concentration and sequences of the input and probe duplex, but did not include the Cy3 fluorophore-quencher pair, which is common in both conditions.

**Figure 4.**
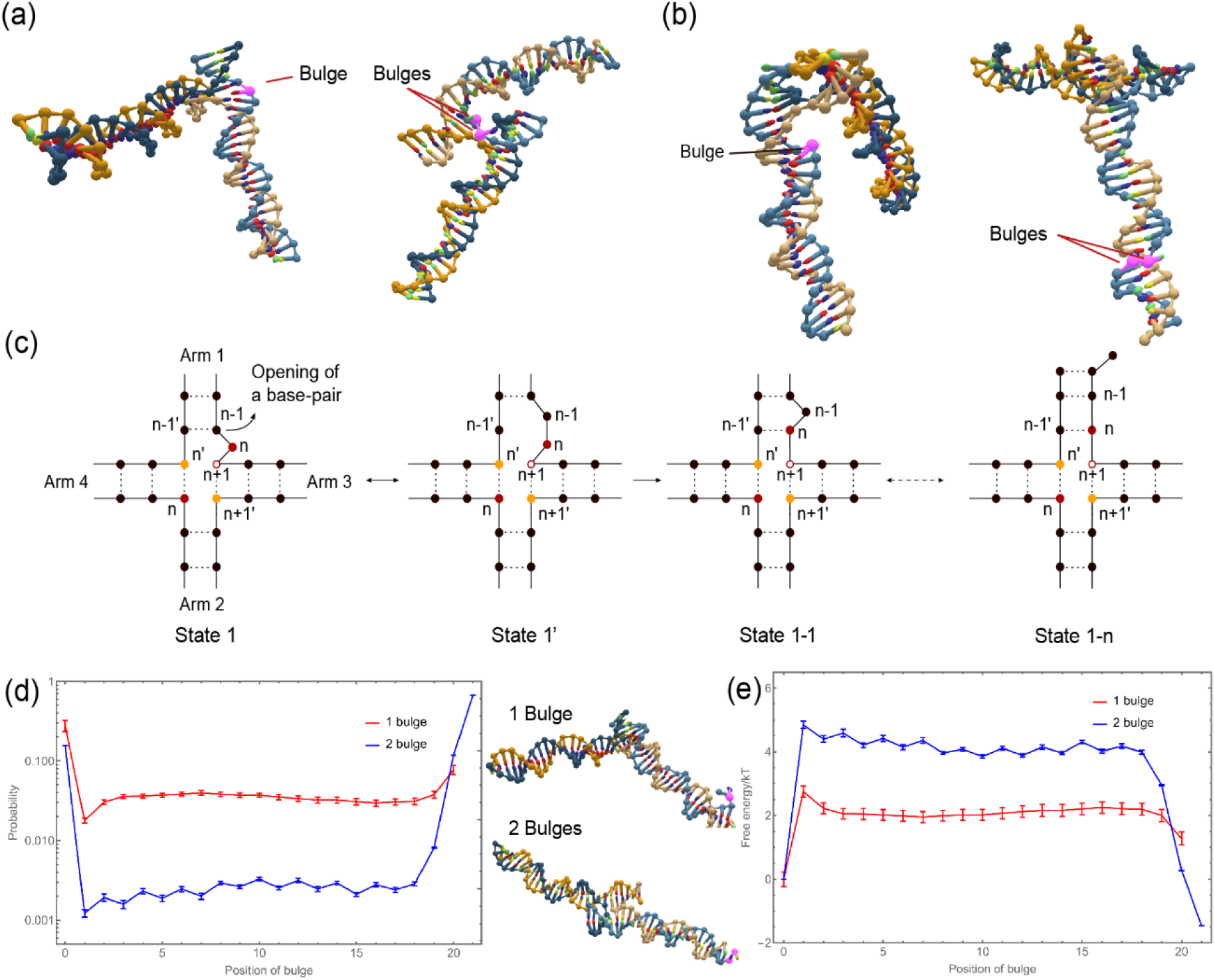
oxDNA predictions of bulge migration in the absence of branch migration. For the 1-bulge and 2-bulge systems, oxDNA simulations were initiated with probe and input bound by toeholds alone. a, b) Representative images of the Holliday junction where the bulges are at the toehold (a), as initiated, and in the middle of branch migration domains (b), as observed later in simulations. c) Proposed states for the diffusion of bulges with a fixed junction. d) Probability distribution of the position of the bulge(s) in the displacement domain of the one-bulge and two-bulge system, given a fixed junction location. Location 0 on the horizontal axis represents the bulge present near the junction, while the rightmost point (20 for the 1 bulge system and 21 for the 2 bulge system) represents the bulge beyond the final position in the displacement domain (where the duplex can enter the frayed state). Representative images of the Holliday junction when the bulges are at location 0 are shown on the right. e) Free energy 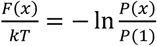 as a function of bulge position *x*, where *P*(*x*) is the probability of finding the bulge at position *x*.

During initial oxDNA simulations, it was observed that the bulge not only participates in the branch migration process but can also diffuse internally along the arm of the input or probe duplex (Figure 4b). While this bulge diffusion does not block the branch migration process, it eliminates the accelerating effect of bulge when the bulge is not in the correct position. For the system with a single bulge, this diffusion is illustrated schematically in Figure 4c. When in state 1, the base pair (*n+1,n+1’*) of arm 3 or (*n-1, n-1’*) of arm 1 can open before the base pair (*n, n’*) of arm 4 (Figure 4c, state 1’), and the bulge can diffuse away from the juction. Due to the free-energy penalty of incorporating a bulge within the interior of a duplex relative to having the extra bases as an overhanging tail, this diffusion is unlikely to happen when the input and probe duplexes are isolated. However, when the duplexes bind to form a junction, those extra bases now destabilize the junction by design. Thus the relative cost of allowing the bulge to diffuse into the interior of the duplex is much lower.

A step of diffusion for the bulge is complete when the new base pair (*n, n+1’*) regenerates the (*n+1*) base of arm 3 as the new bulge (Figure 4c, state 1-1). Similar to the branch migration, this is a reversible step which can proceed to the end of the branch migration domain (Figure 4c, state 1-n).

To explicitly study this bulge diffusion, we ran oxDNA-based simulations while allowing bulge diffusion but prohibiting branch migration. Doing so, we observed the bulge diffusing across the different arms of the duplex (Figure 4d,e). The bulge has two preferred locations: one near the junction and the other at the far end of the duplex arm, where it forms a frayed duplex end rather than an enclosed bulge (Figure 4d, right images). The one bulge system exhibits a slightly higher probability to stay near the junction (0.28 for one bulge and 0.156 for two bulges), potentially explaining the low efficacy of the motif relative to expectations. In the two-bulge system, the bulges have a very high probability to be found at the far end, and a large barrier separating this state from the state with the bulge at the junction. These results suggest a relatively long-lived state with low junction mobility, which may contribute significantly to reducing *k*_bm_. Moreover, the existence of a long-lived low mobility state may explain why the high concentration curves in Figure 3C show anomolously slow convergence to the maximum signal in the final tages of the reaction.

## Discussion

Herein we present an alternative mechanism for four-way branch migration with higher speed limit compared to conventional systems. This increase in speed comes from the introduction of a bulge at the Holliday junction which increases the branch migration rate by more than an order of magnitude. Consequently, this work offers a reaction scheme with improved kinetics applicable to a number of nucleic acid reaction networks that rely on four-way branch migration, including molecular walkers^34^ and DNA actuators^35^.

We noted that introduction of a single bulge enforces a single nucleotide constraint on the displacement domain. Nevertheless, the toehold domains are not subjected to a similar constraint and the use of the 4-way motif, with a larger number of bases required in the toehold than for 3-way strand exchange, increases the discrimination that can be achieved through toeholds alone. Our design remains compatible with the majority of four-way branch migration reaction networks proposed thus far. Indeed, Johnson has pointed out that the ability to use orthogonal branch migration domains in 4-way strand displacement networks is inherently limited anyway^22^.

The presence of bulges increases both *k*_off_ and *k*_bm_ by destabilising the Holliday junction and facilitating branch migrations. In our experiments, this change allowed for faster reactions at higher duplex concentration. Alternatively, *k*_off_ can be supressed by increasing the length of toeholds, allowing for faster reactions at low duplex concentration. Bluge-free systems, by contrast, would run into the same isues with a low speed limit imposed by the low alue of *k*_bm_.

Using oxDNA simulations we were able to partially explain why *k*_bm_ did not increase as much as expected. oxDNA simulations show that the bulge(s) can diffuse along the displacement domain, a process which competes with accelerated junction migration. When the bulge is not at the junction core, the junction migrates as if there is no bulge. Whenever the diffusing junction meets the diffusing bulge during the displacement process, accelerated migration ensues.

We note that the estimated rate of four-way branch migration in the absence of the bulge is an order of magnitude greater than that reported in previous studies^23, 36^. This enhanced rate could be the result of the poly-A/poly-T sequence in the displacement domain, which encourages more rapid branch migration given that A-T base-pairs are weaker than G-C base-pairs^23^. However, we did not see a reduced rate of branch migration in the absence of a bulge when the displacement domain was composed of poly-GT/poly-CA sequences. This may indicate that the difference in branch migration rate results from slight differences in experimental conditions, with the GC-content having only a limited effect.

As mentioned in the introduction, absence of single stranded reaction species makes four-way branch migration ideal for *in vivo* application, due to a reduction in cross-talk and higher stability^29^. However, no method exists for producing multi-stranded nucleic acid duplexes *in vivo*. The recently developed protocol for autonomously generating multi-stranded nucleic acid species^37-38^ can be extended to four-way branch migration reaction schemes. As such, we expect to combine both methods to implement an optimised four-way branch migration scheme *in vivo* as part of future work.

## Data availability

Raw rata, fitting code and oxDNA simulating code are freely available at https://doi.org/10.5281/zenodo.7539760.

## Acknoledgements

W.B. was supported by EPSRC grant EP/P02596X/1. F.S. acknowledges support from the UK Engineering and Physical Sciences Research Council (EPSRC) grant EP/S022856/1 for the Centre for Doctoral Training in BioDesign Engineering. A.S. acknowledges support from the European Research Council (ERC) under the European Union’s Horizon 2020 research and innovation program (Grant agreement No. 851910). T.O. is supported by a Royal Society university research fellowship. G.-B.S gratefully acknowledges support from the UK Royal Academy of Engineering via his Chair in Emerging Technologies (RAEng CiET 1819\5). M.M.S. acknowledges support from the Royal Academy of Engineering Chair in Emerging Technologies award (CiET2021\94).

## Materials and methods

### DNA sequence design

All sequences were designed using NUPACK and ordered from Integrated DNA Technologies (IDT). All strands were ordered with HPLC purification at 100µM in LabReady IDTE buffer (pH 8.0). The sequences used in this work are given in Table 1. Additional clamps (marked with red colour) are placed to ensure correct annealing of displacement domains of the input and the probe complexes.

**Table 1.**
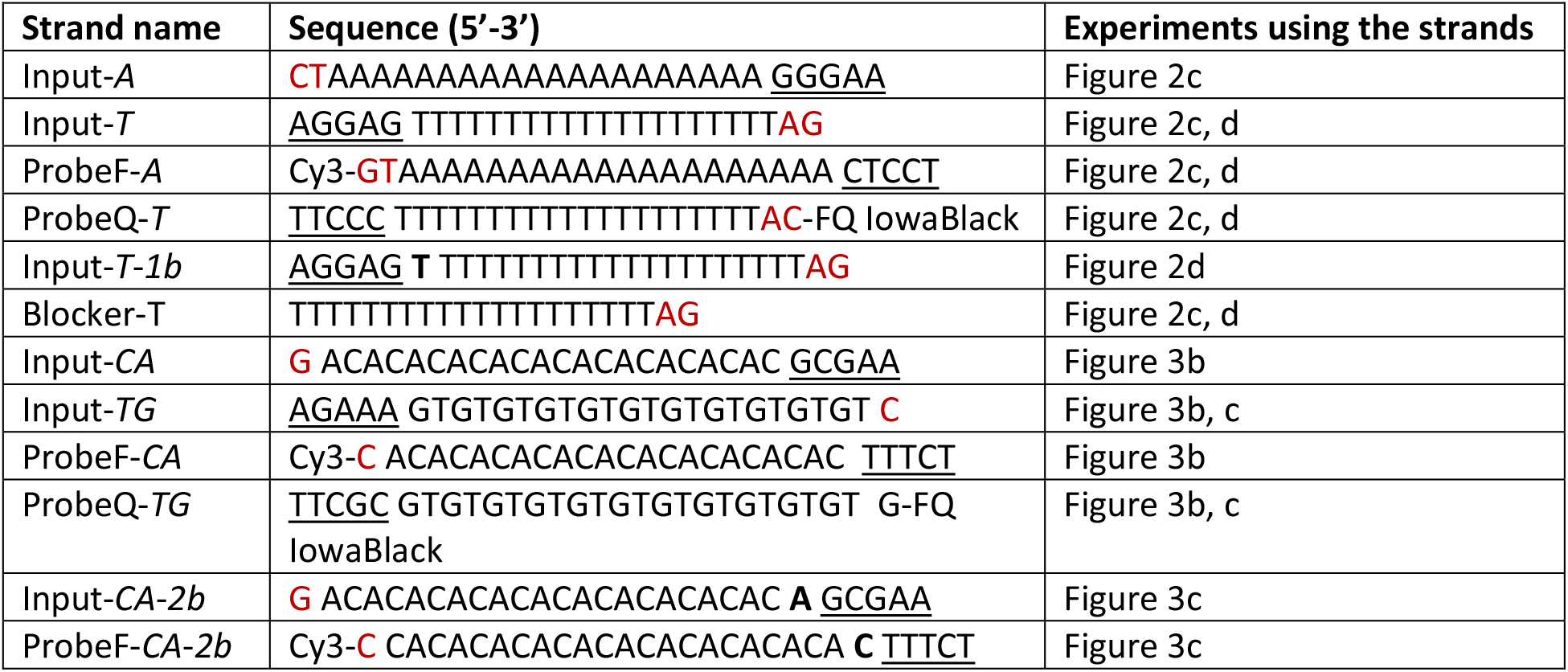
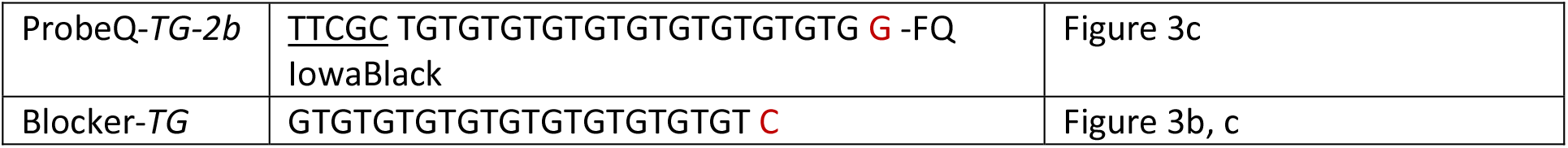
Experimental sequence design. Bulges are shown in bold; toehold is shown with underlines and domains are separated by a space. Cy3 and FQ IowaBlack were used as fluorophore and quencher, respectively. Sequences are given 5’ to 3’.

### DNA duplex annealing

Input and probe duplexes were formed by annealing complementary strands. One of the strands was added at 10% excess in the presence of 20% of blocker strands (Supplemtary note 2). More specifically, 45 µM of poly T (or TG) strand was mixed with 50 µM of poly A (or AC) strands and 10 µM of poly T (or TG) blocker strands in 1X TE buffer with 150 mM of NaCl. Annealing was performed in an Applied Biosystems ProFlex PCR thermal cycler. Strands were annealed by heating to 95°C for 5 minutes and then cooling to 20°C at a rate of 1°C/minute.

### Fluorescence spectroscopy

All fluorescence measurements were performed in a *BMG* Labtech Clariostar microplate reader using *Greiner*’s BioOne flat U-shaped clear bottom 96-well plates. Reactions were performed at a reaction temperature of 25°C. To test different concentrations of probe and input duplexes, probe duplexes were diluted to 6.7 nM and input duplexes were diluted to 40, 200, 400 … nM (4 time of the target concentration) in 1M NaCl + 1X TE buffer. 150 µl of probe duplex solutions were injected to 50 µl of input duplexes at different concentrations. After 2 seconds of mixing by shaking, Cy3 emissions (excitation: 530/20 nm, emission: 580/30 nm) were collected with 570 ms time interval with the first data point at 3.5 seconds. Raw fluorescence values were converted to concentrations by first subtracting background fluorescence from 5 nM of fully quenched probe duplex and then dividing by fluorescence of 5 nM of unquenched Cy3 duplex (ProbeF-*A* / input-*T* or ProbeF-*CA* / Input-*TG* duplexes). 100% of the value corresponds to 5 nM of strand exchanged product.

### Data fitting procedure

All fittings were performed in Python. We described the system using the following ODE model, with rate constants *k*_on_, *k*_off_ and *k*_bm_:

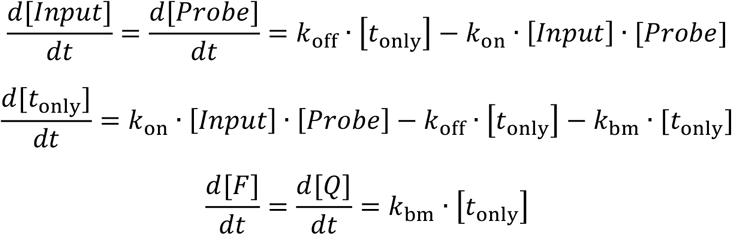

We fit this ODE model to the experimental kinetic data to estimate values for the rate constants *k*_off_ and *k*_bm_. *k*_off_ and *k*_bm_ were allowed to vary between 10^−10^ *s*^−1^ and 10^10^ *s*^−1^, with initial estimates of 10^1^ *s*^−1^. *k*_off_ and *k*_bm_ were estimated as log-transformed values to reduce the overall time to estimation. *k*_on_ was restricted at 10^7^ *M*^−1^ *s*^−1^ to improve the overall fits. We employed a global fitting approach such that a single value of *k*_off_ and *k*_bm_ were estimated for all six concentrations in each experiment (Figure 2c, d and Figure 3b, c). The final mean and standard error reported for each were calculated by jackknife (leave-one-out) estimation. 95% confidence intervals are reported in the original units of measurement.

All data fitting was performed in Python. The Python code for fitting and estimating these rate constant estimates is available at: https://doi.org/10.5281/zenodo.7539760.

### OxDNA simulations

The oxDNA model is a coarse-grained model of DNA that can capture the structural, mechanical, and thermodynamic properties of DNA systems. For further details on how to setup and run the model, and other applications covered with oxDNA, we refer the interested reader to the paper by Segnar et al.^39^

The simulations were performed in a box size of 70 units at 25 °C at 1 M Na. The 4 strands (2 input and 2 probe strands, Table 1) were loaded into the simulation box. Mutually attractive forces were applied (using mutual traps feature in oxDNA) on the complementary bases of input-input and probe-probe strands to bring the system together. This is followed by applying mutually attractive forces on the complementary toeholds (inputA:probeQ-T input:probeF-A for the 1 bulge system and input-CA:probQ-TG and input-TG:ProbeF-CA for the 2 bulge system) to bind all the bases of the toehold. This represents the starting system, which is similar to state 1 in Figure 2(a) for the 1-bulge case, and state 3 in Figure 4(a) for the 2-bulge case.

At any point during the simulation, the bulge can be present at different locations. For the sake of notation, we will hereby refer to the base pairs that are admissible when the bulge is near the junction as the *default* base pairs. When the location of the bulge is somewhere within the arm or at the far end, we will refer to the base pairs as *displaced* base pairs.

3 types of parameters were defined to keep track of the base pairs formed in the 1-bulge system. The first order parameter (OP1) tracks all the default base pairs formed in the respective arms (arm 1 of the one-bulge system) and varies from 0 to 22. The second order parameter (OP2) tracks all the displaced base pairs formed and varies from 0 to 20. The third order parameter (OP3) tracks all the possible base pairs of the other 3 arms. OP1=22, OP2=0 coresponds to the bulge being present at the junction. OP1=2, OP2=20 corresponds to the bulge having diffused to the far end of the duplex but the terminus nucleotides CT in input-A and AG in input-T still forming base pairs. OP1=0, OP2=20 corresponds to the bulge having diffused to the far end of the duplex and the input-A and input-T strands fray at the terminal end.

For the 2-bulge system, an extra order parameter OP4 tracks the base pair at the terminus end of the displacement domain Input-*CA* Input-*TG* (the end farthest from the Holliday junction). OP4=1 means the terminus base pair is fraying and OP4=0 means otherwise. OP1 and OP2 can now take values between 0 and 21. OP1=21, OP2=0, OP3= {0 or 1}^40^ corresponds to the bulge present at the Holliday junction. OP1=0, OP2=21, OP3=1 corresponds to the bulge present at the end of the displacement domain and the terminus base pair is not fraying. OP1=0, OP2=21, OP3=0 corresponds to the bulge present at the end of the displacement domains with the terminus base pair fraying.

Since we are studying the bulge diffusion process and not interested in observing junction migration, we restricted OP3 to retain the maximum number of possible base pairs, preventing junction diffusion. Umbrella sampling^41^ was then used to accelerate opening of base pairs within the arm of interest, which helped to increase the diffusivity of the bulge and allows for sampling of the competition between default base pairs (OP1) and displaced base pairs (OP2). The order parameter files, and the weight files that correspond to the umbrella sampling simulations are available at https://doi.org/10.5281/zenodo.7539760.

It was also observed that simulating the system with 2 bulges was relatively slow. This free-energy barrier that causes this slowdown is visible in Figure 4. The bulges in this case either spend most of their time near the junction or at the ends, spending minimal time inside the displacement domain. To improve the sampling, we perform this simulation in 3 different windows. Each window has an overlap with the other window allowing us to recover the entire landscape using WHAM^42^. The first window restricts the simulations with the condition: OP2<=5 such that the bulge diffuses close to the Holliday junction. The second window samples bulge diffusion for OP1>=4 and OP2>=4. Under this condition, the bulge never reaches either ends of the displacement domain. The third window samples bulge diffusion for OP1<=5. Now, the bulge diffuses near the other end of the displacement domain.The simulations were performed using a VMMC (virtual move Monte Carlo) algorithm. The system was allowed to equilibrate for the first 10^7^ moves. A total runtime of 10^9^ moves for each simulation was used to generate the results presented in Figure 5.

The location of the bulge, as shown in Figure 4(d) and 4(e), is tracked by the value of OP2. Note that for the 2-bulge system to account for the bulge present at the end of displacement domain (OP2=21), we consider both states when terminus base pair is fraying (OP4=0) and when its not fraying (OP4=1).

## Supplementary information

### Supplementary note 1

We assessed the effect of altering the value of *k*_on_ on the estimated value of *k*_bm_. We determine the percentage change in the value of *k*_bm_ compared to the value estimated under the assumption of *k*_on_ = 10^7^ *M*^−1^ *s*^−1^ for each experiment. We identify the value of *k*_on_ which incurs a percentage change of 2.5% in *k*_bm_. For the no bulge system (20nt poly-A/T branch migration domain), the lower bound is *k*_on_ = 10^5.47^and although the exact upper bound was not identified it is greater than *k*_on_ = 10^8.80^, which is already beyond the limit of reasonable *k*_on_ estimates. For the 1-bulge system, the lower bound is *k*_on_ = 10^5.93^ and although the exact upper bound was not identified it is greater than *k*_on_ = 10^8.50^, which again is already beyond the limit of reasonable *k*_on_ estimates. For the no-bulge system (22nt poly-GT/CA branch migration domain), the lower bound is *k*_on_ = 10^5.28^ and although the exact upper bound was not identified it is greater than *k*_on_ = 10^8.80^. For the 2-bulge system, the lower bound is k_on_ = 10^6.20^ and although the exact upper bound was not identified it is greater than *k*_on_ = 10^8.50^. Across all experiments the value of *k*_bm_ differed by less than 2.5% between *k*_on_ = 10^6.2^ and *k*_on_ = 10^8^. We assessed whether these alternative *k*_on_ values equally provided reasonable fits to the experimental data. We observed good fits to the curves at *k*_on_ = 10^8^ across all experimental systems (Figure S1).

**Figure S1.**
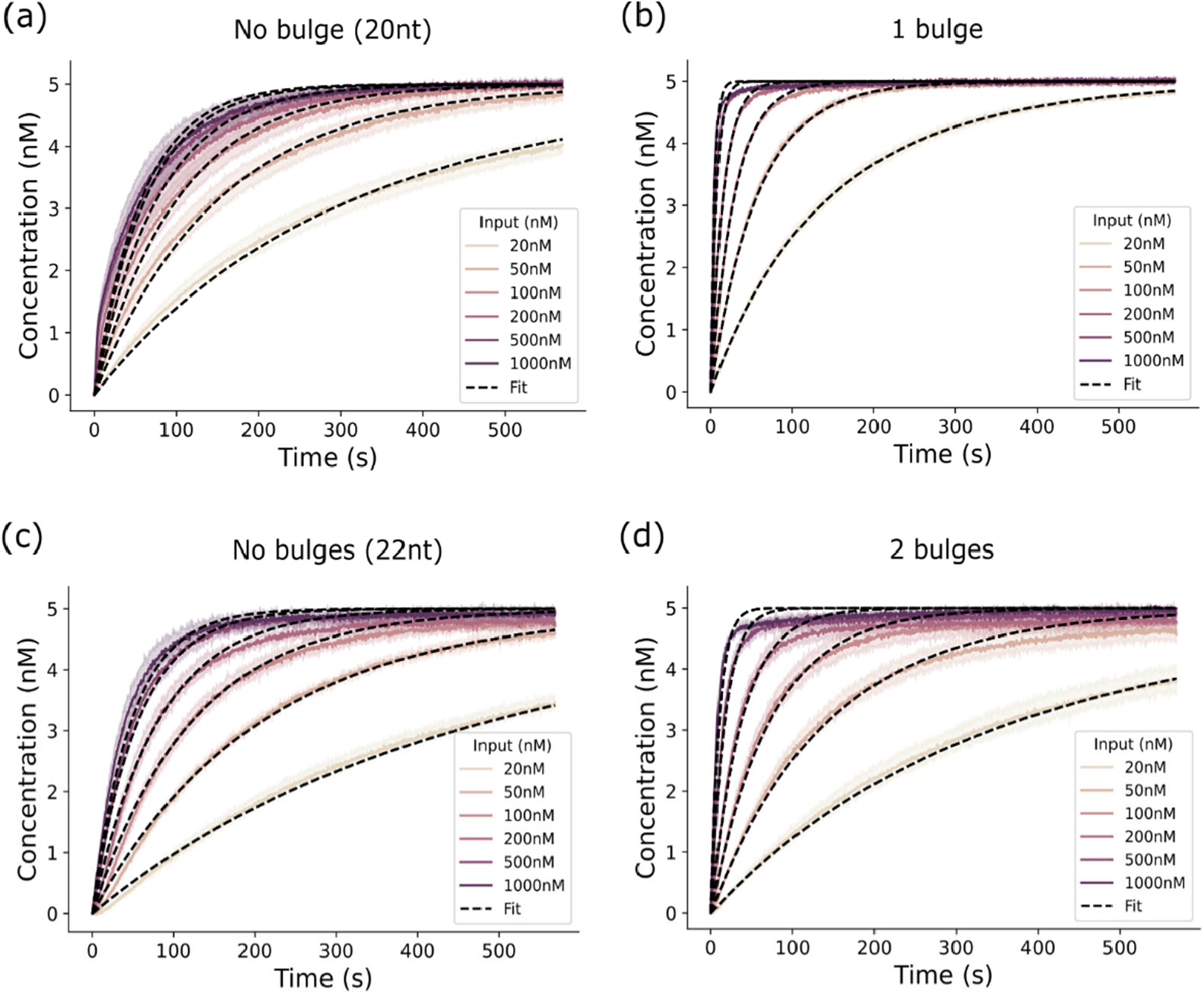
Fits to fluorescence traces under the assumption of *k*_*on*_ = 10^8^. Normalised fluorescent traces from combining 5 nM of probe duplex and 20nM – 1000nM of input duplex in the a) absence or b) presence of 1 bulge and the c) absence or d) presence of two bulges. Black, dashed lines represent fits to the experimental data.

### Supplementary note 2

We performed a series of experiments in the absence of blocker strands. Although we captured the kinetics for the no-bulge system (22nt poly-GT/CA displacement domain) in the absence of blocker strands, we were unable to effectively fit the model to this data. We noted that the estimated rate of branch migration was lower across all experiments in the absence of blocker strands. The estimated values of *k*_off_ and *k*_bm_ for these experiments are reported in Table S1. While the difference in rate in the presence and absence of blocker strands was minimal for the no-bulge system (20nt poly-A/T displacement domain), the rate of the 1-bulge and 2-bulge systems in the presence and absence of blocker strands differed by a factor of 5 and 4, respectively.

**Table S2.** Estimated values of *k*_*off*_ and *k*_bm_ in the absence of blocker strands. Estimated rate constants across a series of four-way branch migration experiments in the absence of blocker strands. 95% confidence for each rate constant estimate is reported.

| Experiment | $k_{off} (s^{-1})$ | 95% CI $k_{off}$ | $k_{bm} (s^{-1})$ | 95% CI $k_{bm}$ |
| --- | --- | --- | --- | --- |
| No bulge | $6.1 \times 10^{-1}$ | [0.56, 0.67] | $1.2 \times 10^{-2}$ | [0.011, 0.013] |
| single bulge | 2.8 | [1.9, 4.2] | $1.1 \times 10^{-1}$ | [0.14, 0.081] |
| two bulges | 5.7 | [4.7, 7.1] | $8.3 \times 10^{-2}$ | [0.070, 0.099] |

